# Cell-in-the-loop pattern formation with optogenetically emulated cell-to-cell signaling

**DOI:** 10.1101/679597

**Authors:** Melinda Liu Perkins, Dirk Benzinger, Murat Arcak, Mustafa Khammash

## Abstract

Designing and implementing synthetic biological pattern formation remains a challenge due to underlying theoretical complexity as well as the difficulty of engineering multicellular networks biochemically. Here, we introduce a “cell-in-the-loop” approach where living cells interact through *in silico* signaling, establishing a new testbed to interrogate theoretical principles when internal cell dynamics are incorporated rather than modeled. We present a theory that offers an easy-to-use test to predict the emergence of contrasting patterns in gene expression among laterally inhibiting cells. Guided by the theory, we experimentally demonstrated spontaneous checkerboard patterning in an optogenetic setup where cell-to-cell signaling was emulated with light inputs calculated *in silico* from real-time gene expression measurements. The scheme successfully produced spontaneous, persistent checkerboard patterns for systems of sixteen patches, in quantitative agreement with theoretical predictions. Our research highlights how tools from dynamical systems theory may inform our understanding of patterning, and illustrates the potential of cell-in-the-loop for engineering synthetic multicellular systems.

## 2 Introduction

Spatial patterning is crucial for the proper functioning of diverse multicellular biological systems from slime molds [1] to developing embryos. The ability to synthetically engineer multicellular patterning will facilitate advances in designing microbial communities [2] [3] [4], creating synthetic biomaterials [5] [6], and programming tissue and organ growth [7] [8] [9] [10], among other applications [11]. While recent efforts to synthetically engineer multicellular patterning have met with success (see [12], [13], [14] for reviews), relatively few of these efforts [15] [16] have been guided by quantitative mathematical theory beyond numerical simulation. In contrast, conventional engineering approaches rely on the predictive power of theory both to design complex systems and to build the intuition necessary to envision new capabilities. Future progress in synthetic multicellular patterning will benefit from a firm understanding of the underlying theoretical principles, as well as scalable, efficient methods for implementing—and validating—these principles in practice.

Gene expression patterning has received much focus in the theoretical literature [17] [18] [19] [20] [21] [22] [23], and is also of particular interest in regenerative medicine, since it is central to the early stages of embryonic development and eventual cell fate determination [7] [24]. There are a number of challenges associated with engineering spontaneous gene expression patterning into biochemical systems, including how to facilitate interaction among cells [25] and achieve spatial precision in the resulting patterns [26] [27] [28]. Even when successful, these implementations are still constrained by time, expense, and the availability of biological parts satisfying parameter requirements [29] [30]. Moreover, it may be difficult to measure or monitor particular system components in real time, which can hinder “debugging” and slow down the design-build-test cycle [31].

While numerical simulation is an important method for efficient prototyping, simulations are only as valid as the models underlying them, and simplifications or faulty assumptions can limit the experimental applicability of simulation results. We propose that future efforts in synthetic patterning would benefit from an intermediate step between pure simulation and full biochemical implementation, which could be used to validate theories or incrementally test synthetic designs before they are fully incorporated into the organism. Inspired by “human-in-the-loop” approaches for engineering systems that must interact with complex, living individuals [32], we propose a “cell-in-the-loop” approach in which physical signaling among cells is substituted with computer-controlled inputs calculated *in silico* from real-time measurements of gene expression. Cell-in-the-loop, by incorporating live cells into the “simulation”, eliminates the need to make assumptions about individual cell behavior during dynamic evolution, while retaining flexibility in testing parameters that remain under computational control. These benefits are particularly essential for patterning systems, in which the large number of interacting cells can make detailed simulations prohibitive or impossible.

We implemented cell-in-the-loop using optogenetics, which have been shown to afford excellent spatiotemporal precision in applications including feedback control [33] [34] [35] [36] and oscillatory synchronization [37]. We engineered *Saccharomyces cerevisiae* to respond to blue light [38] by increasing gene expression as measured by a fast-acting fluorescent reporter [39]. We used an optogenetic platform capable of targeting individual cells independently of each other [36], such that the light input to any given cell could be calculated based on the gene expression levels of other cells that were interacting with the target cell. Both the network architecture (which cells interacted with which) as well as the exact form of interaction were programmed into the computer, allowing us to precisely modulate system parameters related to cell-to-cell signaling.

We adapted a general theory for pattern emergence in large-scale lateral inhibition systems [40] [41] to inform our designs and predict steady-state outcomes. Specifically, we programmed a computational *signaling relation* to emulate mutual inhibition among groups of cells and varied the strength of the inhibition by tuning a single digital bifurcation parameter. Once the network architecture and signaling relation were defined, inputs to cells were calculated solely based on measurements of those cells without any further external control, creating a self-contained dynamical system. Using this setup, we visualized gene expression levels of real cells by the brightness of square patches on a virtual grid (Fig. 1). Depending on the value of the bifurcation parameter, the system was expected to produce a homogeneous non-patterning outcome or generate contrasting “checkerboard” patterns in which neighboring patches alternated between expressing high and low levels of gene. We showed that checkerboard patterns could indeed emerge spontaneously in our system and that patterning outcomes were persistent across the last hour of the experiment. Although stochastic effects dominated the precise outcomes of individual experiments, the theory qualitatively predicted pattern emergence and quantitatively predicted average patch brightness across a range of parameter values. Our results demonstrate the utility of a cell-in-the-loop approach for designing and evaluating systems of interacting cells, as well as probing the limits of deterministic theory in the face of stochastic influence.

**Figure 1:**
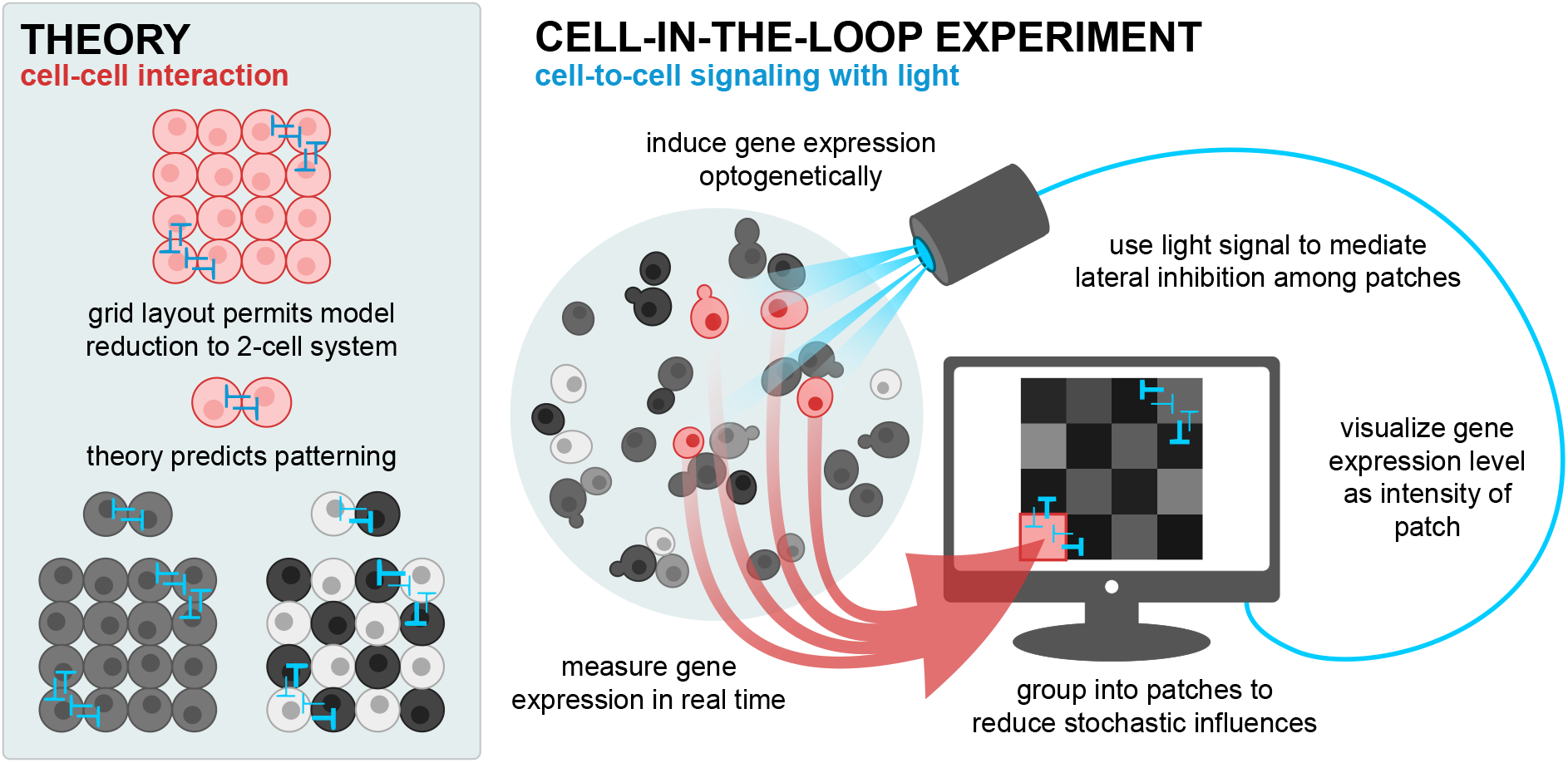
Spontaneous checkerboard patterning with optogenetically emulated cell-to-cell signaling. Optogenetically responsive cells signal to each other through computer-controlled light inputs that vary in intensity based on the gene expression levels of other cells. We enact lateral inhibition according to the theory in Section 3.1 that predicts when cells will spontaneously separate into two classes of high and low gene expression. In all figures, red denotes *in vivo* and blue denotes *in silico* components.

## 3 Results

### 3.1 Theory predicts checkerboard patterning using a test for bistability

We developed theory to predict the emergence of stable contrasting patterns in deterministic systems of laterally inhibiting cells [40] [41]. Here, we adapt the theory to the present optogenetic implementation. We emphasize how our system was decomposed into *in vivo* and *in silico* components, each of which corresponds to a particular element in the theory, and how this correspondence enables empirical measurement and experimental design.

Consider a system of *N* isogenic cells signaling to each other. Suppose we measure for each cell a scalar output such as fluorescence that correlates positively with gene expression level and is designated by *w*_*i*_ for the *i*th cell. The input *u*_*i*_ to a cell affects output levels with an empirically characterizable dose response, which describes the steady-state level of *w*_*i*_ for a constant-in-time input. In our setup, the input *u*_*i*_ is light, and increasing input intensity increases gene expression. This portion of the theory represents the *in vivo* component of the system.

To synthesize *u*_*i*_, we first average the measured gene expression, *w*_*j*_, over all cells *j* signaling to cell *i*, and denote this average as *v*_*i*_. We then set the input to the *i*th cell to *u*_*i*_ = *h*(*v*_*i*_), where *h*(*⋅*) is the signaling relation programmed into the computer. To enact mutual inhibition, increasing gene expression in one cell must decrease gene expression in neighboring cells. Therefore, since higher-intensity light induces higher gene expression, we select *h*(*⋅*) to be decreasing.

We chose a grid layout with periodic boundary conditions in which each cell signals four other cells reciprocally. This layout satisfies all assumptions discussed in the box entitled “Summary of theory”, therefore we can predict contrasting patterning in a full system of *N* cells based on the bistability of an equivalent 2-cell system. If the 2-cell system is monostable, then both cells express the same level of gene, and the *N*-cell system also has a stable state in which all cells express the same level of gene. Inversely, if the 2-cell system is bistable, then one of the stable states corresponds to one cell expressing high levels of gene and the other, low, and the other stable state corresponds to the opposite situation. In this case, two stable, contrasting steady-state patterns also exist for the *N*-cell system; that is, one subset of the *N* cells expresses identically high levels of gene, and the remaining cells express identically low levels of gene (or vice versa). Contrasting steady states can be visualized as checkerboards in which neighbors alternate between high and low. The 2-cell system can be assessed for bistability using a standard technique illustrated in Fig. 2 and described in Section S1.1.

**Figure 2:**
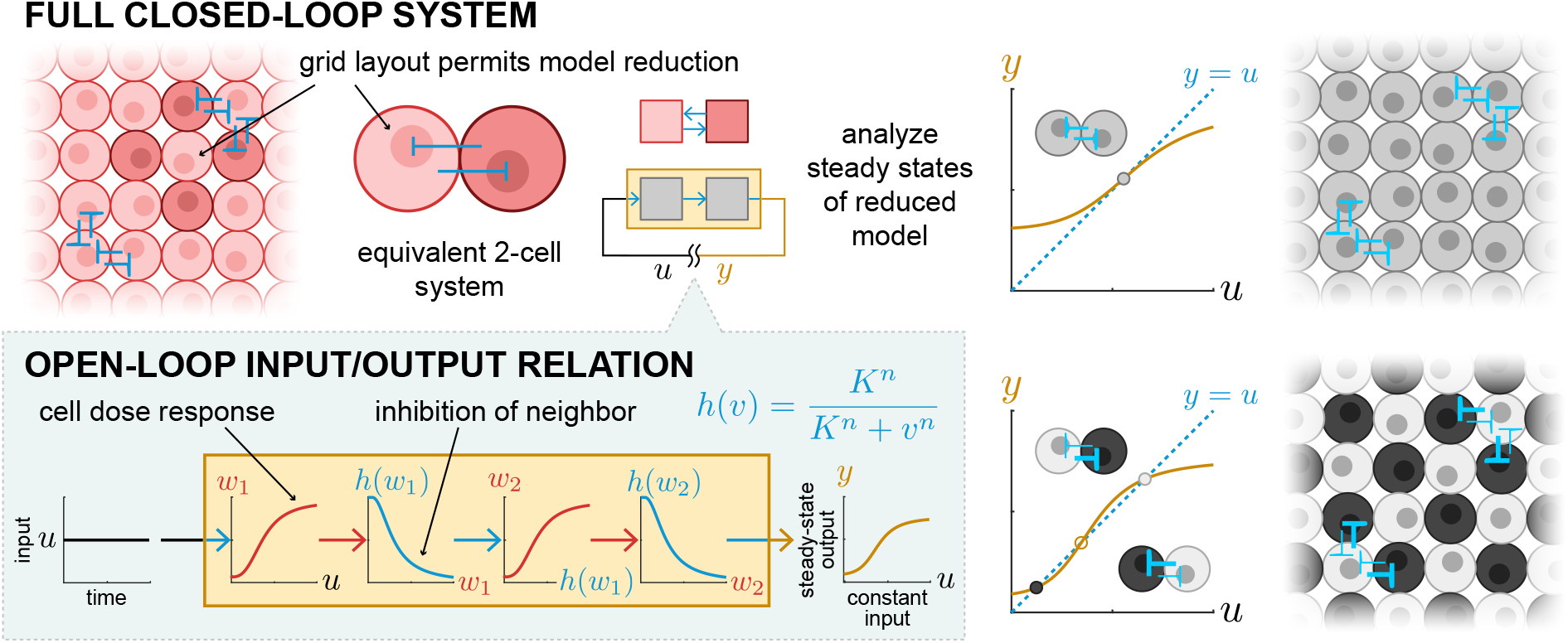
Spontaneous contrasting patterning in a large lateral inhibition system can be predicted from bistability in a 2-cell system. The bistability of a 2-cell system can be used to evaluate contrasting patterning in a full network of isogenic, mutually inhibiting cells arranged in a grid with periodic boundary conditions. Bistability of the corresponding 2-cell system implies contrasting patterns exist in the full system (see box entitled “Summary of theory”). Bistability can be assessed through an analytical test in which the closed-loop dynamical system is “opened” into an input/output system by breaking the feedback loop. If a constant-in-time input to the open-loop system produces a steady-state output value equal to the input value, then that value is a steady state for the corresponding closed-loop system.

To apply the theory, we must know (1) the dose response, in our case *in vivo* gene expression levels under varying intensities of light; and (2) the form of the signaling relation, here programmed *in silico*. Thus, to carry out lateral inhibition experiments, we needed to measure an empirical dose response of cells to light, and define the computational signaling relation controlling light inputs such that intensity was inversely related to the responsiveness of cells interacting with the target.

### Summary of theory

Consider a system of *N* identical cells modeled as single-input, single-output dynamical systems. Biochemical concentrations 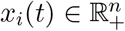 in the *i*th cell evolve according to

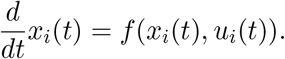

Each cell has output *w*_*i*_(*t*) = *g*(*x*_*i*_(*t*)) *∈* ℝ_+_ and input *u*_*i*_(*t*) *∈* ℝ_+_. Let the vector 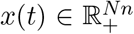 be the vertical concatenation of the vectors *x*_*i*_(*t*) for all *N* cells, and similarly for 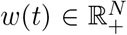 and *u*(*t*) *∈* ℝ^*N*^. We assume each cell has a *static input-output characteristic T* (*⋅*), that is, if a cell is given constant-in-time input 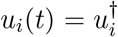, it will reach a globally asymptotically stable hyperbolic equilibrium 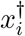 solving 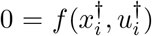 with output 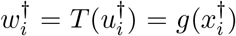. We assume *T* (*⋅*) is bounded and increasing, meaning that increasing the input increases the output. In our setup, the static input-output characteristic corresponds to the empirically measured dose response.

Suppose the outputs of cells are connected to the inputs of other cells, forming a network. We capture information about which cells signal to which by way of the interconnectivity matrix 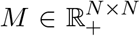 with entries [*M*]_*ij*_ = 0 if cell *j* does not signal to cell *i* and [*M*]_*ij*_ *>* 0 otherwise, with the value [*M*]_*ij*_ indicating the strength of signaling. We require that the sum over all entries in a row equal the same constant, *µ ∈* ℝ_+_, regardless of the row, i.e., Σ_*j*_[*M*]_*ij*_= *μ* for all *i*. In our setup each cell receives signals from four other cells with equal weights 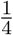, therefore *μ* = 1. Defining *v*(*t*) = *M w*(*t*), we model lateral inhibition by letting the input to cell *i* be given by *u*_*i*_(*t*) = *h*(*v*_*i*_(*t*)), where *h*(*⋅*): ℝ_+_ *→* ℝ_+_ is bounded and decreasing.

#### Model reduction theorem

*Let* 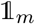 represent the length-m column vector of all ones, and similarly for 0_*m*_. *If there exists a matrix* 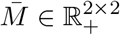 *such that*

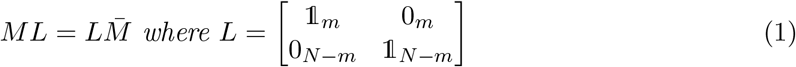

for some indexing of cells, then 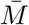 is an interconnectivity matrix for an equivalent 2-cell system whose steady-state solutions correspond to steady states of the N-cell system with interconnectivity matrix M. In other words, if 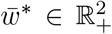 is the output corresponding to a steady-state solution to the 2-cell system, then 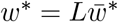 *is a steady-state output to the N-cell system*.□

Note that the cells indexed 1 through *m* take on steady-state output values 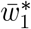 while those indexed *m* + 1 through *N* take on steady-state output values 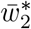. Condition (1) is satisfied when cells can be grouped into two subsets within which nodes are interchangeable; that is, reindexing nodes within a subset will not change *M* [41].

When

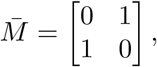

the steady states of the 2-cell system are determined graphically from the fixed points of *h*(*T*(*h*(*T*(⋅)))), as shown in Fig. 2 and explained in Section S1.1. For the reduced 2-cell system the graphical test also ensures stability of the points corresponding to the lower/upper intersections in Fig. 2, and instability of the point corresponding to the middle intersection, when the cellular dynamics are *monotone* in the input/output sense [42]. The stability properties established graphically for the 2-cell system are preserved in the full N-cell system when additional assumptions hold. Our setup satisfies one such assumption from [40], which stipulates cells within a subset not signal to each other. Thus, if 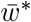 is the output corresponding to a stable state in a bistable 2-cell system, then in the N-cell system, cells in one subset have higher output than cells in the other subset. If the cells belonging to different subsets are spatially interlaced or alternating, then the high/low dichotomy produces a spatially contrasting pattern such as a checkerboard.

### 3.2 Empirical characterization informs computational parameter choice

We combined the blue light-inducible VP-EL222 expression system [43] [38] with a fast-acting nuclear translocation reporter (dPSTR) [39] to control and measure gene expression in *Saccharomyces cerevisiae*. In the dark, constitutively expressed red fluorescent protein (RFP) fused to the synthetic bZip domain SZ2 [44] is equally distributed between nucleus and cytoplasm due to passive diffusion through the nuclear membrane. Under exposure to blue light, VP-EL222 molecules dimerize and bind the cognate promoter to activate expression of a protein comprising two nuclear localization signals (NLS) and SZ1 [44]. This protein then forms a heterodimer with the RFP reporter, thereby localizing fluorescence in the nucleus. We quantitated the degree of nuclear localization (nuclear localization score) as the difference between mean cytoplasmic and mean nuclear fluorescence normalized to the mean fluorescence across the entire cell. In principle, the score is 0 if cells are not at all responding (there is no nuclear localization) and positive otherwise (Fig. 3a,b).

**Figure 3:**
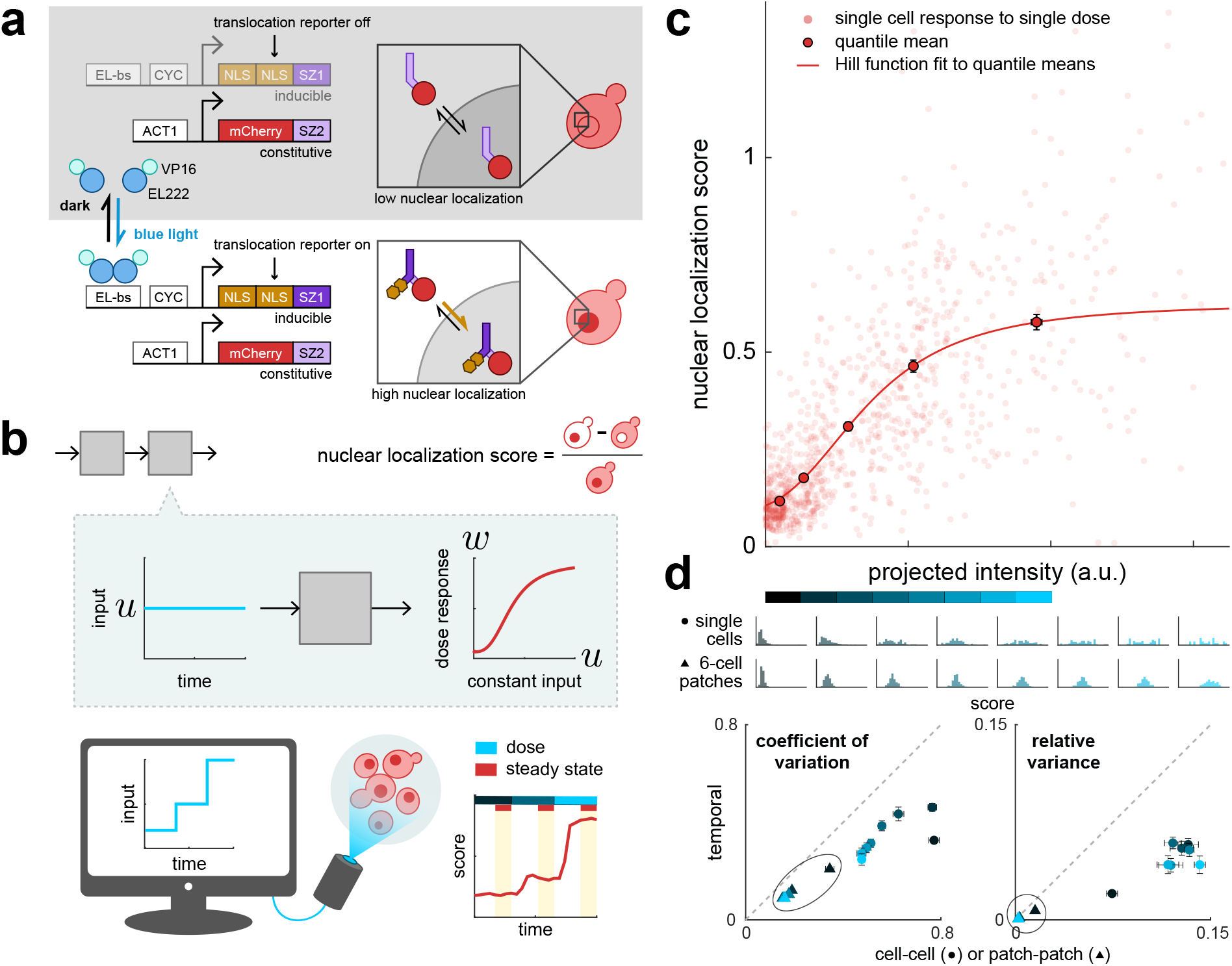
Experimental characterization of cellular dose response curves shows population-level gradedness. (a) **Blue light induces gene expression in optogenetically responsive *Saccharomyces cerevisiae* via the VP-EL222 system [36], as measured by a nuclear translocation reporter (dPSTR) [39].** Cells constitutively express RFP fused to SZ2. In the dark, RFP is distributed equally between nucleus and cytoplasm due to diffusion. Blue light induces VP-EL222 dimerization, which activates expression of an NLS fused to SZ1. SZ1 binds SZ2, such that the NLS promotes localization of RFP in the nucleus, increasing nuclear fluorescence relative to cytoplasmic fluorescence. (b) For a single dose, individual cells were illuminated for 80 min with a constant intensity of light. Single-cell responses to a single intensity were calculated as the average score over the last 40 min. Pictured in the schematic are average input intensities and responses for approx. 700 cells for 3 consecutive doses. (c) **Although individual cell responses vary, on the population level nuclear localization is graded with respect to the intensity of the light input.** Circles without outlines correspond to responses of single cells to single doses across 3 experiments. The final dose response was determined as a Hill function fit to quantile means (solid outline). Error bars are standard error. (d) **Grouping cells into patches reduces both cell-cell and temporal variability.** Scores were binned by projected intensity (blue shaded bar) to generate histograms of score distributions for given input levels. The bin cutoff is twice the maximum projected intensity used in the final patterning experiments. Histograms for 6-cell patches were generated for each bin by bootstrapping from individual cell scores within that bin. For an ergodic process, we would expect cell-cell and temporal variation to be equal (gray dotted line); here, responses appear to be more variable from cell to cell than for single cells across time. Grouping cells into patches of 6 across which response scores are averaged (triangles) reduces the magnitude of difference between cell-cell or patch-patch and temporal variability.

We characterized the dose response of individual cells to constant, targeted blue light exposure (Fig. 3b,c). During these and subsequent patterning experiments, cells were grown in a monolayer under the microscope and automatically imaged, segmented, tracked, and scored by a software pipeline integrating open-source software YouScope [45] with custom Matlab® scripts. On average, cells exhibited a graded response to light intensity well described by a Hill function (Fig. 3c). Variability from cell to cell was greater than for individual cells across time, perhaps owing to variation in cell cycle state [46]. As the theory is deterministic, for patterning experiments we ultimately substituted single cells with computationally defined patches of 4 or 6 cells, with the patch response determined as the average response of the constituent cells. Generating score distributions for such patches by bootstrapping from the single-cell dose response data shows reduced temporal and patch-patch variability as well as reduced difference between temporal and patch-patch variability relative to the single-cell case (Figs. 3d, S1.3).

Based on the range of cellular response scores, we defined the signaling relation *h*(*⋅*) for use in patterning experiments, which determined the light input administered to a patch as a function of the average scores of neighboring patches at each time step. We chose an inhibiting Hill function with fixed Hill coefficient *n* = 2 and a single free parameter *K* with smaller values corresponding to sharper inhibition. We combined the empirical dose response with the computational signaling relation to generate theoretical predictions for the mono- or bistability of a 2-cell lateral inhibition system as *K* was varied between 0 and 1, corresponding to non-patterning or patterning outcomes in a full system.

### 3.3 Cell-in-the-loop lateral inhibition spontaneously generates checkerboard patterning

We ran a series of patterning experiments emulating lateral inhibition. Cells were randomly assigned to patches such that cells belonging to the same patches were not necessarily neighbors in physical space, thereby reducing spurious correlations that might arise from spatially dependent factors other than the targeted light input. Once assigned, cells remained in the same patch throughout the duration of an experiment. Patches were arranged to neighbor each other in “virtual space” as visualized on a checkerboard (Fig. 4a).

**Figure 4:**
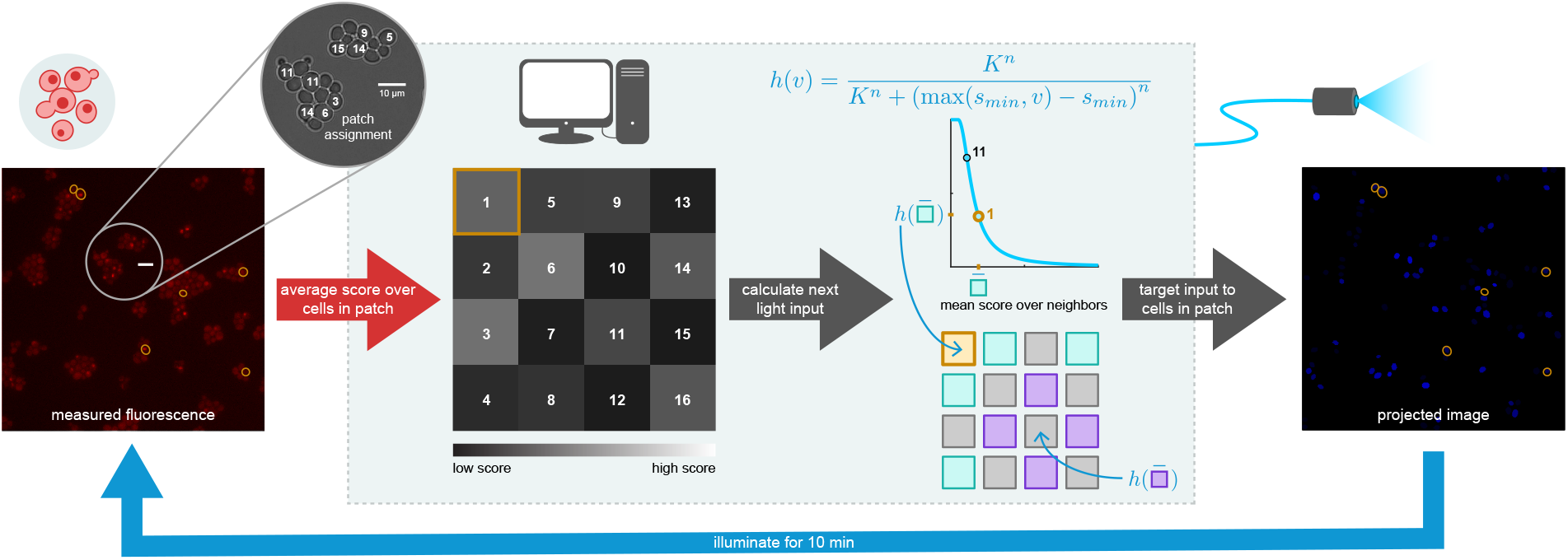
Schematic of automated workflow for patterning experiments. Cells growing in a monolayer were placed under the microscope, which was connected to a computer system that controlled the camera and blue light projection system. Cells were segmented and tracked in brightfield images and scored in fluorescence images (here, false-colored red). After the first image was acquired, cells were randomly assigned to patches to which they belonged for the remainder of the experiment. Grid visualizes scores at a single time step, with each square representing one patch of cells. Brightness corresponds to the average score of the constituent cells in the corresponding patch. The signaling relation, defined before experiment start, determined the input light intensity administered to patches for the next time step (10 min) based on the scores of neighboring patches to the north, south, east, and west, with periodic boundary conditions. Cells in the same patch received the same input intensity targeted individually to each cell, as shown in the projected image.

During patterning experiments, cells were imaged and inputs adjusted every 10 min. We tested systems of 16 patches with 6 cells per patch for four values of *K* between 0.1 and 1. Spontaneous patterning was always achieved in the *K* = 0.1 case and never in the *K* = 1 case, with mixed results for *K* = 0.2, 0.3, near one of the theoretically predicted critical points (Fig. S1.4a). Sample time traces at *K* = 0.1 and *K* = 1 show, respectively, the gradual deviation in score between sets of alternating patches that characterizes a contrasting pattern, or a rapid adoption of a non-patterning state. Visualizing the checkerboard at individual time points or averaged over the last hour clearly depicts the distinction between the two cases (Fig. 5a).

**Figure 5:**
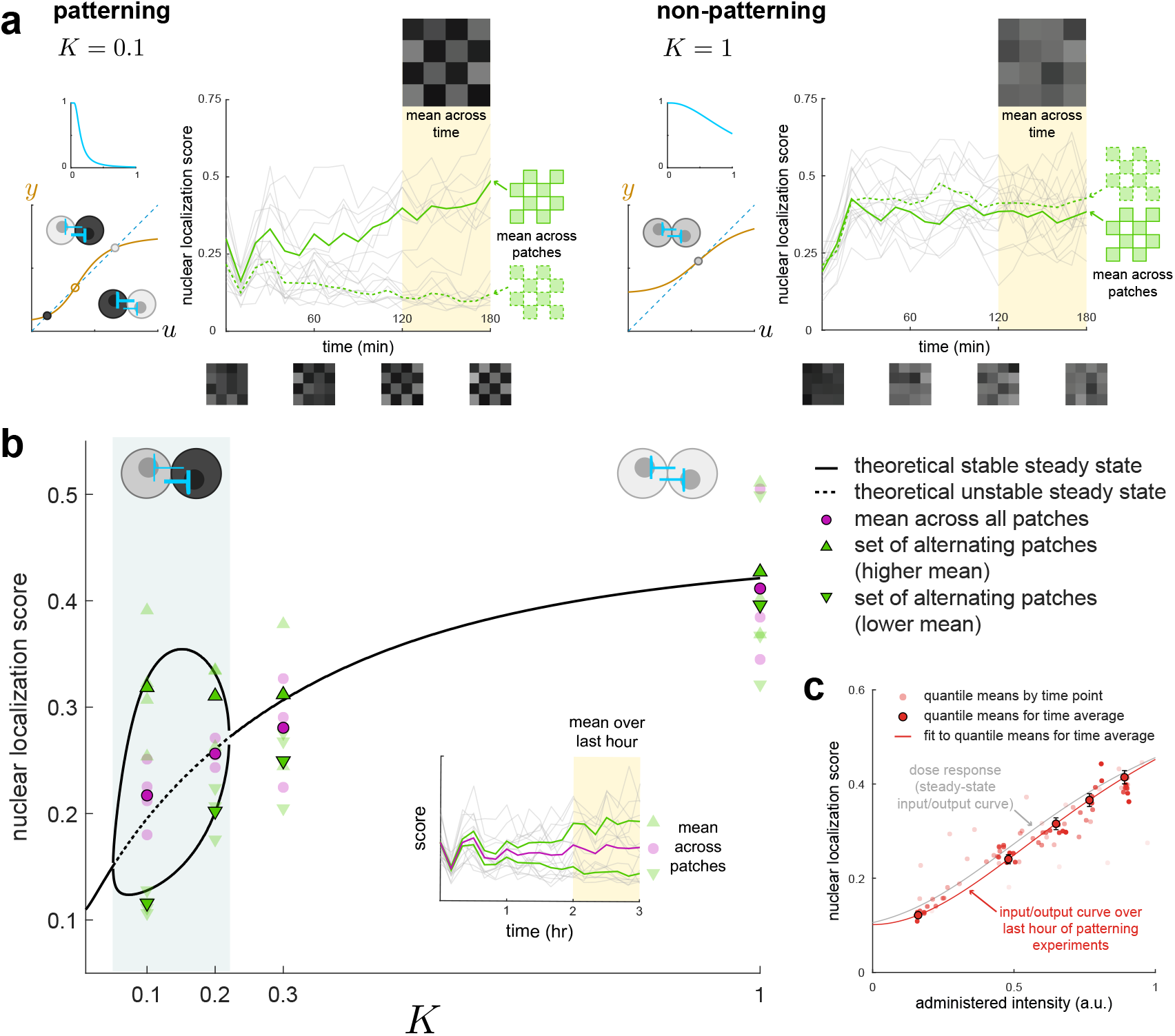
Theory quantitatively predicts spontaneous patterning and patch intensity during cell-in-the-loop experiments. **(a) Sample time traces show emergence of a contrasting pattern (*K* = 0.1) or convergence to a non-patterning state** (*K* = 1). Gray lines correspond to score traces for individual patches; green lines indicate the mean scores of sets of alternating patches. Checkerboards visualize scores at single time points (bottom) or averaged over the last hour (top). **(b) Theory quantitatively recapitulates experimental results for mean patch response score.** Black lines denote theoretical steady-state points as a function of the bifurcation parameter *K*, with solid indicating stable and dashed, unstable. All points are averaged over the last hour. Faded points correspond to individual experiments; solid outlines are averages across experiments (*N* = 4 for *K* = 0.1, *N* = 3 for *K* = 0.2, 0.3, 1). Magenta circular points are averages over all patches. Green points are averages over sets of alternating patches, with upward- and downward-facing triangles denoting the higher and lower of the two means respectively. Stochasticity in the experimental system introduced contrast between the average means of sets even in regions that were deterministically suggested to be monostable. As predicted, the contrast level (difference between means of sets of alternating patches) was higher for lower *K*. Experiment-to-experiment variation in overall brightness (score averaged across all patches) was greater for higher *K*, an effect that cannot be accounted for in a purely deterministic theory. **(c) The 16-patch system with 6 cells per patch converges to a steady state by about 2 hr into patterning experiments.** Solid outline, score values averaged over the last hour for individual patches and split into quantiles by administered intensities show decent agreement with the empirical steady-state dose response curve. Error bars are standard error. Circles without outlines are quantiles at individual time points pooled from *N* = 13 experiments (darker red at later times). For comparison, the dose response curve fit to empirical data is plotted in gray.

When averaged over the last hour and across experiments, the contrast level (mean scores of sets of alternating patches) was quantitatively well predicted by theory in the bistable region and the overall brightness (mean score across all patches) was well predicted in the monostable region (Fig. 5b). When considering individual experiments, the variability in overall brightness increased with increasing *K*. In the predicted monostable cases, stochasticity also introduced a difference between the means over alternating patches, though statistical analyses confirm that the difference was indistinguishable from random (Table S1.4). Taken together, these results suggest that the deterministic theory calibrated to population averages is an excellent quantitative predictor for mean system behavior across time, patches (cells), and experiments, while at the same time even small amounts of cell-cell variability and temporal stochasticity may cause a given experiment to deviate considerably from quantitative forecasts.

Because our setup allowed us to monitor both gene expression levels and cell signaling levels, we were able to assess convergence to (quasi-)steady state by comparing instantaneous input-output curves (patch score vs. administered intensity) to the steady-state dose response (Fig. 5c). Specifically, since the time for cells to converge to steady state under exposure to light of constant intensity (40 min) was longer than the time between changes to input intensity during experiments (10 min), the instantaneous input-output curve during a patterning experiment would only match the empirical dose response curve if the administered intensity remained relatively constant for several frames before a given time point—i.e., if there was little temporal variability for at least 40 min preceding the frame. Directly plotting the temporal variability in administered input to individual patches does indeed reveal a decrease from the first to the last experimental hour regardless of *K* value (Fig. S1.5a).

Lastly, we examined the effect of patch number on patterning outcomes through four experiments with 36 patches, 4 cells per patch, and *K* = 0.1. None of the experiments spontaneously achieved a checkerboard pattern across the whole board in 3 hr, although a control experiment preinduced with the pattern showed that it was indeed persistent (Fig. S1.7). The input/output curve did not approach the empirical dose response (Fig. S1.6) and temporal variability in administered intensity was the same during the first and third experimental hour (Fig. S1.5b), further supporting the conclusion that the system never reached steady state. Interestingly, one experiment produced two checkerboards in opposite corners that persisted throughout the last experimental hour, but were inverted relative to each other and did not resolve before the end of the experiment (Fig. S1.4b). Other experiments also exhibited transient local patterning, although to a lesser degree. The local patterning and the increased convergence time are consequences of the fact that a 36-patch system admits a much larger space of possible configurations than a 16-patch system. Although variability in 4-cell patches was only modestly larger than in 6-cell patches (Fig. S1.3), the stochasticity may also have contributed to a longer convergence time. These and related challenges will require further investigation in future efforts to synthetically generate gene expression patterns with single-cell granularity.

## 4 Discussion

In this work, we employed cell-in-the-loop, a closed-loop, hybrid *in vivo*/*in silico* approach, to validate a theory for spontaneous gene expression patterning among laterally inhibiting cells. We engineered *S. cerevisiae* to respond optogenetically to light inputs, then emulated cell-to-cell signaling in real time by modulating the intensity of light inputs to cells based on real-time measurements of gene expression. The theory made accurate quantitative predictions for average steady-state patterning outcomes across a range of parameters. Increasing system size—by increasing the space of possible dynamic behaviors—diminished the probability of achieving global patterning on short timescales in the absence of initial or external bias. Further theroetical research should explicitly incorporate cell-cell variability and temporal stochasticity in order to improve our understanding of variation in individual experimental outcomes, patterning robustness, and the link between individual-level and population-level behavior.

Prior work using optogenetics to generate persistent spatial patterns in living cells has focused on reproducing [47] [48] [49] or processing [50] pre-existing images projected by the light input. In comparison to these studies, light in our system does not *a priori* encode a pattern to which the cells conform; rather, light acts as a virtual signal transmitted from cell to cell. The input intensity is determined by cellular responses that are in turn influenced by the received intensity, establishing a closed-loop relationship independent of external control. That similar patterns are ultimately observed in both the cellular responses and the optogenetic inputs arises as a consequence of their mutual dependence.

Depending on the application, cell-in-the-loop offers benefits over purely biochemical or purely computational approaches. First, it reduces the number of components that must be engineered into cells. We integrated a single optogenetically induced promoter and a single reporter, and were able to modulate patterning outcomes simply by reprogramming the computer. In this way, we circumvented issues associated with synthetic cell-to-cell signaling, including parameter matching and crosstalk [51] [28], and alleviated complications such as burden [52] [53] that arise from integrating complex networks into cells [29] [31]. We were also able to achieve spatiotemporal control over the whole population of interacting individuals and probe stochasticity at a finer level than would be attainable with a conventional biochemical implementation.

Compared to a computer simulation, cell-in-the-loop makes no assumptions about cell behavior or the form of biological noise, since the cells themselves are incorporated into the system. Although we used this setup to test the validity of a theoretical principle, one could also envision testing the accuracy of a full model for cell-to-cell signaling by simulating a proposed physical mechanism of interaction, then comparing the outcome of such a system to the outcome of a purely physical system. Cell-in-the-loop also allows one to track system components that might otherwise be inaccessible or difficult to measure. For example, we were able to monitor the levels of both gene expression and virtual signal simultaneously, which could be difficult to achieve in a solely biochemical setup.

Once established, a cell-in-the-loop system could couple with more complex cellular processes to achieve real-world results. One could also envision using cell-in-the-loop as a rapid prototyping platform or “stepping stone” to a fully biochemical implementation. In this paradigm, one would begin with minimally engineered cells and then sequentially replace *in silico* components with biochemical ones, testing at each stage whether the remaining portions of the network ought to be modified in structure or value before the next component is incorporated into the cell. Such an approach could also reveal shortcomings in proposed designs; for example, in our setup, 36-patch systems failed to produce spontaneous patterns on our experimental timescale even with “perfect” deterministic signaling, suggesting that it could be challenging to achieve large-scale lateral inhibition patterning biochemically in a similar context. Thus, the addition of cell-in-the-loop to the biological engineering process could greatly decrease the time, expense, and effort required to develop synthetic multicellular systems for an increasingly rich and promising array of applications.

## 5 Methods

### 5.1 Plasmid and yeast strain construction

*E. coli* TOP10 cells (Invitrogen) were used for plasmid cloning and propagation. The dPSTR reporter plasmid (pDB161) contains the coding sequences of UbiY-2xSV40NLS-SynZip1 [39], expressed from an EL222-responsive promoter (p5xBS-CYC180) [38], and mCherry-SynZip2 [39], expressed from the constitutive *ACT1* promoter. It was constructed by first replacing the promoter p*RPL24A* in the plasmid pDA183 [39] by p*ACT1* using SacI-XbaI cut sites and subsequently replacing the promoter p*STL1* by P5xBS-CYC180 using PCR and SapI-based Golden Gate cloning [54].

The yeast strain used in this study (DBY165) was constructed by transforming the PacI digested plasmid pDB161 into DBY41 [38], a strain with BY4741 background expressing VP-EL222 [43] from the *ACT1* promoter. The transformation was performed using the standard lithium acetate method [55].

### 5.2 Culture preparation

Cells were grown at 30°C in synthetic medium (SD) consisting of 2% glucose, low fluorescence yeast nitrogen base (Formedium), pH 5.8, 5 g/l ammonium sulfate, and complete supplement of amino acids and nucleotides. Cultures were started from plate, diluted, and maintained at OD_600_ *<* 1.5 between 24 and 32 hr before an experiment. For each experiment, between 3 and 5 mL of cell culture were centrifuged at 20°C, 3000 RCF for 6 min and enough supernatant was removed to achieve an approximate OD_600_ of 4 after resuspension. Cells were then immediately placed on agarose pads, prepared according to the procedure in Section S1.2.1.

### 5.3 Imaging

Images were taken under a Nikon Ti-Eclipse inverted microscope (Nikon Instruments) with a 40x oil-immersion objective (MRH01401, Nikon AG, Egg, Switzerland), pE-100 bright-field light source (CoolLED, UK), and CMOS camera ORCA-Flash4.0 (Hamamatsu Photonic, Solothurn, Switzerland) water-cooled with a refrigerated bath circulator (A25 Refrigerated Circulator, Thermo Scientific). The temperature was maintained at 30°C by an opaque environmental box (Life Imaging Services, Switzerland), and a dark cloth was additionally placed over the microscope to fully shield cell samples from external light. Experiments were conducted with a diffusor and a green interference filter placed in the bright-field light path, with the Nikon Perfect Focus System (+/-5 AU) enabled. Fluorescence images were acquired using a Spectra X Light Engine fluorescence excitation light source (Lumencor, Beaverton, USA), filter cube with excitation filter 565/24 nm, emission filter 620/52 nm, and beam splitter HC BS 585 (AHF Analysetechnik AG, Tübingen, Germany). The final fluorescence images used for analysis were maximum projections across z-stacks of 5 images spanning 0.6 *μ*m.

During experiments, the microscope was operated by the open-source software YouScope [45]. Cell segmentation and tracking were performed on brightfield images using software tools developed by [36] based on [56] and [57]. For each cell, the mean fluorescence in the nucleus, cytoplasm, and across the entire cell were automatically calculated using custom Matlab (MathWorks) scripts following the procedure in Section S1.2.2.

### 5.4 Light-delivery system

Optogenetic inputs were delivered to cells using the setup developed in [36], in which images generated on the computer are projected by a digital mirror device through a system of lenses that focuses the light onto a microscope slide. Two neutral density filters (Thorlabs, 25 mm absorptive, optical densities 0.5 and 1.3) were placed serially to achieve a total density of 1.8.

To ensure light mapped properly from the projector to the cell, images were modified prior to projection in order to map pixels on the DMD to pixels in the camera images. The mapping was determined through the procedure outlined in [36], Fig. S6B. The procedure was performed immediately before experiment start on an area of the agarose pad unoccupied by cells.

We observed that there was a sigmoidal relationship between the administered illumination intensity and the measured illumination intensity in the projection images, and also that images of uniform intensity did not evenly illuminate the sample plane. To compensate for these effects, we modified images before projection to linearize the administered-to-measured intensity relationship and also to reduce the intensity of overilluminated regions to match the level attained by underilluminated regions. Details of the procedure are available in Section S1.2.5.

Custom Matlab® code was used for manually calibrating projector intensity before experiment start (Section S1.2.5) and for automatically carrying out experiments.

### 5.5 Dose response

The theory relies on a deterministic dose response curve in which the expected steady-state response score of a cell increases as a function of constant input intensity. We performed a series of dose response experiments to verify that these conditions held for our yeast strain and then calculated a dose response curve from the average response of cells to varying measured projected intensities.

Cells tended to respond much more strongly and unpredictably to the first administered input than to later inputs regardless of the intensity of the first input. Therefore, before administering any doses, all cells on the dish were illuminated for 10 min with uniform, middling intensity light, then left in the dark for 20 min to allow the response to decay. Multiple doses were then administered in immediate succession to cells on the plate. For a single dose, cells were illuminated with individually targeted light with constant administered intensity per cell. Cells were imaged every 10 min. Cells that were not successfully segmented and tracked at all sampled time points in a dose were discarded. The steady-state response of a single cell was calculated as the average of the scores from 40 to 80 min under illumination.

The final dose response curve was fit to data aggregated from three experiments. Individual cell responses were binned by projected intensity into 5 quantiles. The final dose response curve in the form of a leaky activating Hill function

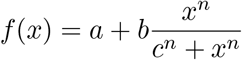

was fit to the mean score values within each quantile vs. the mean measured projected intensity within that quantile.

### 5.6 Patterning experiments

Before experiment start, all cells on the plate were illuminated for 10 min with uniform, middling intensity light, then left in the dark for 10 min such that responses would not fully decay, allowing for some initial variation in nuclear localization score. Experiments lasted 3 hr, during which cells were imaged and their inputs adjusted every 10 min. Preliminary experiments confirmed that cells were alive and responsive up to 6 hr after placement under the microscope, although final experiments were constrained to 3 hr to ensure cells remained in a monolayer.

#### 5.6.1 Patch construction

Cells were randomly assigned to groups with a fixed number of cells per group. Each group corresponded to a single computationally defined “patch”. The score for a patch was calculated as the average of the scores of the cells comprising the patch. By “administering an input to a patch”, we mean the cells in that patch were individually targeted with the same administered input. Most experiments were performed with 16 patches of 6 cells per patch. Higher-dimensional experiments were performed with 36 patches of 4 cells per patch.

Patches were arranged in “virtual space” into a square grid where each patch was connected to (interacted with) each of the patches to the north, south, east, and west, with periodic (wraparound) boundary conditions such that each patch interacted with four other patches (its “neighbors”). Note that, because cells were randomly assigned to patches, cells to neighboring patches in virtual space were not necessarily adjacent in real space.

#### 5.6.2 Signaling relation

Every imaging period (10 min), the input to each patch was adjusted according to the signaling relation, which was chosen to be a Hill function with a minor computational adjustment to better utilize the available range of illumination intensities. Specifically, the relation was given by

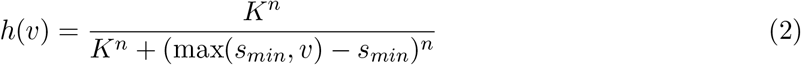

where *v* was the average score across all four neighbors of a patch, *n* was fixed at 2, and the minimum score *s*_*min*_:= 0.05 was the cutoff to deliver maximum illumination intensity. The bifurcation parameter *K* varied by experiment, with higher *K* corresponding to weaker repression.

In particular, let 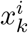 be the score of the *i*th patch in the *k*th frame, and 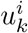 be the input to the patch between the *k*th and (*k* + 1)th frames. Then 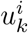 was calculated as

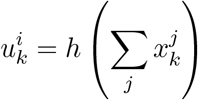

where the summation is taken over the neighbors of patch *i* and *L*(*⋅*) is as given in (2).

## 6 Acknowledgments

The authors would like to thank Marc Rul*⋅*lan for his help with the setup and guidance throughout the project, as well as Brian M. Lang for his advice on statistics and general linear modeling for the dose response data. This project received funding from Air Force Office of Scientific Research (AFOSR) grant FA9550-14-1-0089 and the European Research Council (ERC) under the European Unions Horizon 2020 research and innovation programme (CyberGenetics; grant agreement 743269).

## S1 Supplementary material

### S1.1 Graphical test for bistability

Consider a 2-cell system of mutual inhibition with dynamics (and notation) as given in Section 3.1, in which *u*_2_(*t*) = *h*(*v*_2_(*t*)) = *h*(*w*_1_(*t*)) and similarly *u*_1_(*t*) = *h*(*v*_1_(*t*)) = *h*(*w*_2_(*t*)). Now break the loop such that *u*_1_(*t*) becomes the input and *y*(*t*) = *h*(*w*_2_(*t*)) the output of a the *open-loop system*. The static input-output characteristic for the cascade, given by 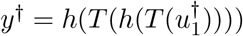, is increasing. The points where 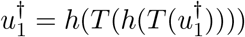 (i.e., intersections of the static input-output characteristic with a line of slope 1) are the steady states of the corresponding *closed-loop system* in which *u*_1_(*t*) = *y*(*t*) (and we still have *u*_2_(*t*) = *h*(*w*_1_(*t*))). We will designate values at such intersections by superscript asterisks (e.g., 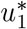). When cell dynamics are monotone, then the steady state corresponding to 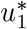 is stable if the slope of 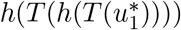 is greater than 1 and unstable if it is less than one. In particular for our setup, this implies that if there is one intersection, the closed-loop system is monostable, and if there are three intersections, the system is bistable, with the homogeneous solution 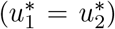 being unstable and the other two stable points corresponding to 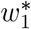 high, 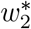 low, and vice versa [58].

### S1.2 Detailed methods

#### S1.2.1 Agarose pad preparation

**Figure S1.1:**
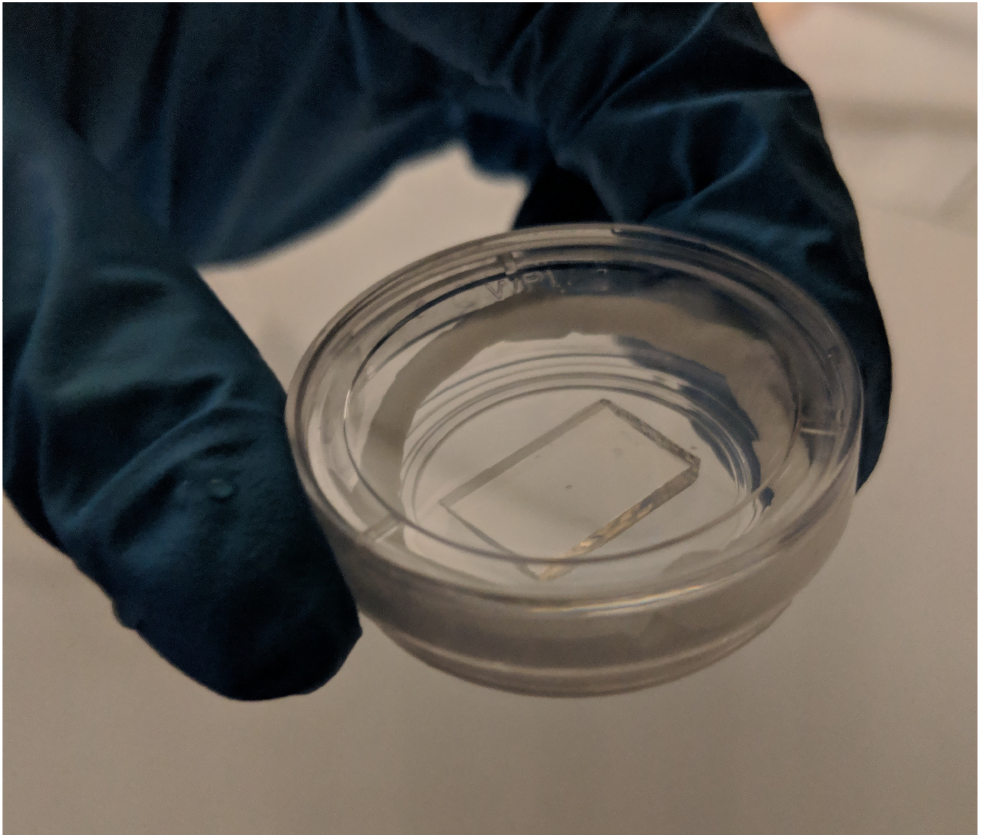
Example agarose pad used in experiments.

Two 2-slide-tall stacks of microscope slides were placed 1 cm apart parallel to each other on top of a single microscope slide. 70 *μ*L of 2% agarose (UltraPure^TM^Agarose, Invitrogen) and SD medium solution were pipetted between the two stacks. A square 18 mm *×* 18 mm cover slip was gently placed on the top. The pad was solidified for 1 hr. Immediately before placement under the microscope, the stacks and cover slip were removed and the ends of the pad were sliced off with a scalpel such that the final pad was about 15 mm *×* 15 mm and level across the top. 3 *μ*L of cell suspension were pipetted in increments of 1 *μ*L onto three separate areas of the pad to ensure that at least one would have the correct density for use in the experiment. The pad was overturned into a circular tissue culture dish with cover glass bottom (35 mm FluoroDish^TM^, World Precision Instruments) lined inside with a strip of damp paper towel to maintain humidity throughout the course of the experiment. The dish was closed and sealed with a strip of parafilm, then placed in the microscope’s environmental box (Life Imaging Services, Switzerland). Cells were allowed to settle for 30 min before experiment start.

#### S1.2.2 Scoring

**Figure S1.2:**
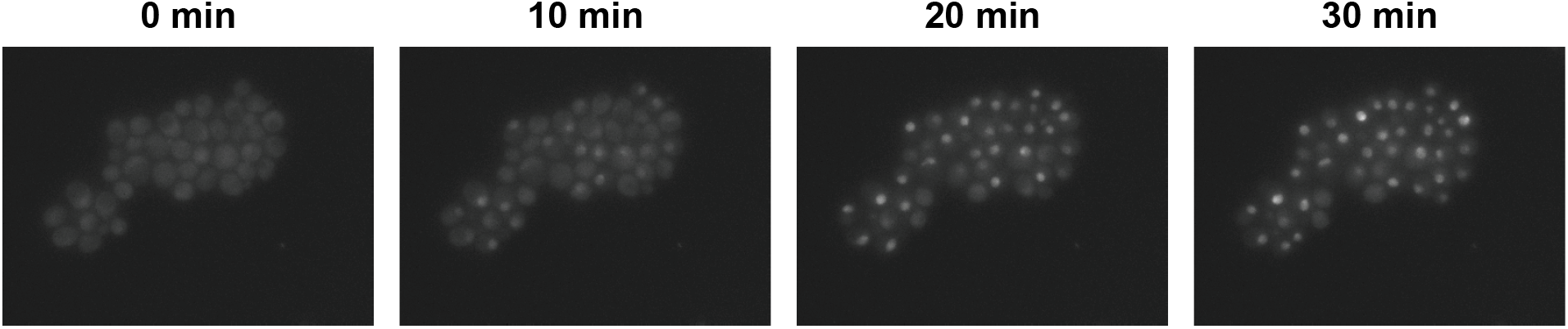
Example time lapse of maximum projection fluorescence images for cells under constant illumination.

We assessed induction of gene expression using a fast-acting, nuclear translocation-based reporting system [39], where higher responses corresponded to greater fluorescence in the cell nucleus. Accordingly, we defined the scoring scheme as

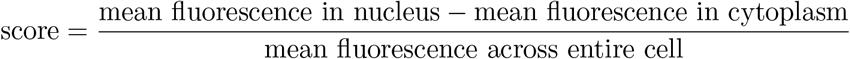

such that a score of 0 indicates no difference between nuclear and cytoplasmic fluorescence.

Our experiments required us to calculate the response score automatically at each time step. The score for each automatically segmented cell was calculated for an approximated location of the nucleus, determined as follows: First, an image was formed by taking a box around the segmented cell in the maximum projection fluorescence image and converting the non-cell pixels to black. A black border 5 pixels wide was added around the image. Then the image was blurred with a Gaussian filter of standard deviation 2 pixels and the brightest pixel in the blurred image was located. This point was considered to be the center of the nucleus. We noted by observation that the nucleus was almost always 5 pixels in radius, therefore the pixels falling within a circle of radius 5 pixels around the center were presumed to belong to the nucleus and were subsequently used in the calculations of the mean fluorescence. All remaining pixels belonging to the cell were considered to belong to the cytoplasm. The score was then calculated as indicated above.

#### S1.2.3 Segmentation/tracking errors

The automated imaging pipeline occasionally failed to identify cells in particular frames. In the majority of cases the system was able to recover the cell within one or two frames. In dose response experiments, cells that were not tracked for the entirety of a dose were discarded. During patterning experiments, cells that were not tracked in a frame did not receive input for the following 10 min, and scores for their corresponding patches were calculated as averages over the scores of the remaining cells. In no experiments were all cells in a patch simultaneously dropped in the same frame.

**Table S1.2:** Fraction of all measurements (one measurement = a single cell in a single frame) that were dropped in each experiment.

|  | K | Fraction Dropped Measurements |
| --- | --- | --- |
| **16-patch** | 0.1 | 0.041666667 |
|  | 0.1 | 0.029605263 |
|  | 0.1 | 0.049342105 |
|  | 0.1 | 0.026864035 |
|  | 0.2 | 0.038377193 |
|  | 0.2 | 0.038377193 |
|  | 0.2 | 0.091557018 |
|  | 0.3 | 0.10252193 |
|  | 0.3 | 0.097039474 |
|  | 0.3 | 0.064144737 |
|  | 1 | 0.029057018 |
|  | 1 | 0.04879386 |
|  | 1 | 0.090460526 |
|  | K | Fraction Dropped Measurements |
| **36-patch** | 0.1 | 0.094298246 |
|  | 0.1 | 0.041666667 |
|  | 0.1 | 0.119152047 |
|  | 0.1 | 0.148391813 |

#### S1.2.4 Preliminary dose response experiments verify gradedness and independence of dose history

In order for the theory to apply, we needed to verify (a) that the magnitude of cell response increased with received light intensity; and (b) that the dose response would be independent of dose history for the duration of the final patterning experiments. By “dose history”, we refer to the number, intensity, and order of doses administered to cells prior to a particular time of interest.

The following experiments were performed with an ND filter of optical density 2 (Thorlabs, 25 mm absorptive). Cells were illuminated for 10 min with uniform light, then left in the dark for 20 min before doses were administered. Cells were imaged every 10 min. Doses were administered for 40 min and the administered intensity of a dose varied for different collections of cells on the same plate simultaneously. Hereafter, we use “dose response experiment” to refer to a particular collection of cells on the same plate receiving the same series of administered doses. Two dose response experiments were conducted simultaneously per plate, with four or five doses per experiment. For a single time point, the measured projected intensity received by a cell was calculated as the average of the mean measured projected intensity across all pixels occupied by the cell.

A series of general linear models were fit to the data for each cell. The natural log of the score (response variable) was treated as a function of the agarose pad, frame, dose number (ordered by time of appearance during experiment), frames since dose start, current measured illumination intensity, and the measured illumination intensity for all frames up to the minimum experimental duration (16 frames) before the current frame. Intensities for time points before the start of the experiment were set to 0. An analysis of deviance for the full model (Table S1.2) suggests that the illumination history past the current illumination contributes little to the current score.

Furthermore, we determined by observation that cells had reached a quasi-steady state response before 30 min, and found that fitting a general linear model to steady-state times only (30 min and 40 min into a dose) greatly diminishes the importance of plate, frame, dose number, and frames since dose start (Table S1.2). Moreover, a reduced model for the steady-state score that includes only the current intensity has an AIC substantially similar to that of the full steady-state model (2675 for the reduced vs. 2622 for the full). Together, these analyses suggest that it is reasonable to consider steady-state dose response as a function of current illumination intensity alone.

**Table S1.2:** Results for a general linear model of cell response score across four experiments at all points in time. ImX indicates the measured projected intensity preceding the current by X frames (0 is current frame). The AIC for this model is 9745. As expected, the time since dose start contributes a large drop in deviance, as cells did not instantly settle to a new steady state when the intensity was changed.

| Variable | Df | Deviance | Resid. Df | Resid. Dev | Pr(>Chi) |
| --- | --- | --- | --- | --- | --- |
| NULL | NA | NA | 4117 | 3905.828561 | NA |
| Im0 | 1 | 1051.353764 | 4116 | 2854.474798 | 0 |
| Im1 | 1 | 148.2219005 | 4115 | 2706.252897 | 1.65E-53 |
| Im2 | 1 | 0.229287964 | 4114 | 2706.023609 | 0.544725097 |
| Im3 | 1 | 4.35808067 | 4113 | 2701.665528 | 0.008275942 |
| Im4 | 1 | 1.320218071 | 4112 | 2700.34531 | 0.146119572 |
| Im5 | 1 | 0.074542198 | 4111 | 2700.270768 | 0.729833304 |
| Im6 | 1 | 0.240410136 | 4110 | 2700.030358 | 0.535126523 |
| Im7 | 1 | 1.161222828 | 4109 | 2698.869135 | 0.17286582 |
| Im8 | 1 | 0.187482186 | 4108 | 2698.681653 | 0.583904657 |
| Im9 | 1 | 2.286447549 | 4107 | 2696.395205 | 0.055792838 |
| Im10 | 1 | 0.249079692 | 4106 | 2696.146126 | 0.527855487 |
| Im11 | 1 | 0.596308281 | 4105 | 2695.549817 | 0.328685196 |
| Im12 | 1 | 0.142415919 | 4104 | 2695.407402 | 0.633115034 |
| Im13 | 1 | 0.323652973 | 4103 | 2695.083749 | 0.471767645 |
| Im14 | 1 | 0.836572299 | 4102 | 2694.247176 | 0.24730139 |
| Im15 | 1 | 1.060493657 | 4101 | 2693.186683 | 0.192713497 |
| Im16 | 1 | 3.170352656 | 4100 | 2690.01633 | 0.024309053 |
| FramesSinceDoseStart | 1 | 106.9288382 | 4099 | 2583.087492 | 4.29E-39 |
| Pad | 1 | 13.86651829 | 4098 | 2569.220973 | 2.47E-06 |
| Frame | 1 | 0.002190109 | 4097 | 2569.218783 | 0.952796489 |
| DoseNumber | 1 | 9.159326761 | 4096 | 2560.059457 | 0.00012912 |

**Table S1.2:** Results for a general linear model of cell response score across four experiments for steady-state frames (30 and 40 min after dose start) only. ImX indicates the measured projected intensity preceding the current by X frames (0 is current frame). Note that the frames since dose start, pad, frame, and dose number are less significant in this model relative to the full model in S1.2. The AIC for this model is 2622.

| Variable | Df | Deviance | Resid. Df | Resid. Dev | Pr(>Chi) |
| --- | --- | --- | --- | --- | --- |
| NULL | NA | NA | 1185 | 1140.234564 | NA |
| Im0 | 1 | 481.0998049 | 1184 | 659.1347596 | 1.03E-201 |
| Im1 | 1 | 15.65834255 | 1183 | 643.476417 | 4.58E-08 |
| Im2 | 1 | 0.206888356 | 1182 | 643.2695287 | 0.529734222 |
| Im3 | 1 | 1.351828641 | 1181 | 641.9177 | 0.108199518 |
| Im4 | 1 | 1.058581525 | 1180 | 640.8591185 | 0.155179456 |
| Im5 | 1 | 0.658939118 | 1179 | 640.2001794 | 0.262075464 |
| Im6 | 1 | 0.111574008 | 1178 | 640.0886054 | 0.644450512 |
| Im7 | 1 | 0.009561578 | 1177 | 640.0790438 | 0.892536317 |
| Im8 | 1 | 2.783572744 | 1176 | 637.2954711 | 0.021164596 |
| Im9 | 1 | 1.374292916 | 1175 | 635.9211782 | 0.105311673 |
| Im10 | 1 | 3.334521299 | 1174 | 632.5866569 | 0.011640495 |
| Im11 | 1 | 1.633214394 | 1173 | 630.9534425 | 0.077459177 |
| Im12 | 1 | 0.519193207 | 1172 | 630.4342493 | 0.319493668 |
| Im13 | 1 | 0.000278015 | 1171 | 630.4339712 | 0.981621402 |
| Im14 | 1 | 3.592110318 | 1170 | 626.8418609 | 0.008831956 |
| Im15 | 1 | 2.201528614 | 1169 | 624.6403323 | 0.040371525 |
| Im16 | 1 | 3.868935675 | 1168 | 620.7713966 | 0.006577405 |
| FramesSinceDoseStart | 1 | 4.202474492 | 1167 | 616.5689221 | 0.004622429 |
| Pad | 1 | 0.000989428 | 1166 | 616.5679327 | 0.965336552 |
| Frame | 1 | 6.225817085 | 1165 | 610.3421156 | 0.000566296 |
| DoseNumber | 0 | 0 | 1165 | 610.3421156 | NA |

#### S1.2.5 Ensuring uniform illumination intensity

We calculated our intensity corrections based on the following model: If *u* is an *N* × *N* input image (normalized to [0, 1]) and *u*_*r*_ is that image magnified to size *M* × *M*, then the measured intensity *y* (normalized to [0, 1]) is an *M* × *M*-dimensional image given by

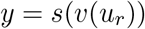

where *s*(*⋅*) is a sigmoidal function applied identically to each pixel, and *v*(*⋅*) is a function that varies by pixel to represent the uneven illumination intensity. This suggests that, to achieve a desired *y*_*d*_, an input image should be calculated as

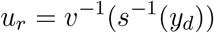

(which can be appropriately resized to obtain *u*).

*s*^−1^(*⋅*) was already determined from previous experiments with the setup [36]. This correction was calculated once, as it did not appear to change between experiments. *v*^*−1*^(*⋅*) was calculated before each experiment by sampling the measured intensity at a number of points spaced across the sample plane when the administered intensity was maximized. To do so, one hundred circles arranged in a grid were projected one at a time onto the slide at maximum administered intensity and the reflected images for each circle were measured. The mean measured intensity of all the pixels in a single circle was treated as a sample of the “true” illumination intensity at the point on the sample plane corresponding to the center of the circle, such that a 2D quadratic polynomial surface could be fit to all mean circle intensities in order to interpolate the illumination intensity at all points on the sample plane. These sample points were normalized to the maximum and a second 2D quadratic polynomial surface was fit to the inverse sigmoid of these normalized sample points. A target intensity value was chosen and the surface fit was renormalized relative to this intensity value to obtain a matrix *V* representing the factor by which bright areas were overilluminated relative to areas of underillumination, such that the matrix 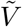 consisting of the element-by-element inverse of 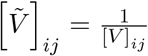 was normalized to [0, 1]. Subsequently, inputs *u*_*r*_were calculated to achieve the desired output *y*_*d*_ as

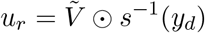

where ⊙ indicates element-by-element multiplication.

**Figure S1.3:**
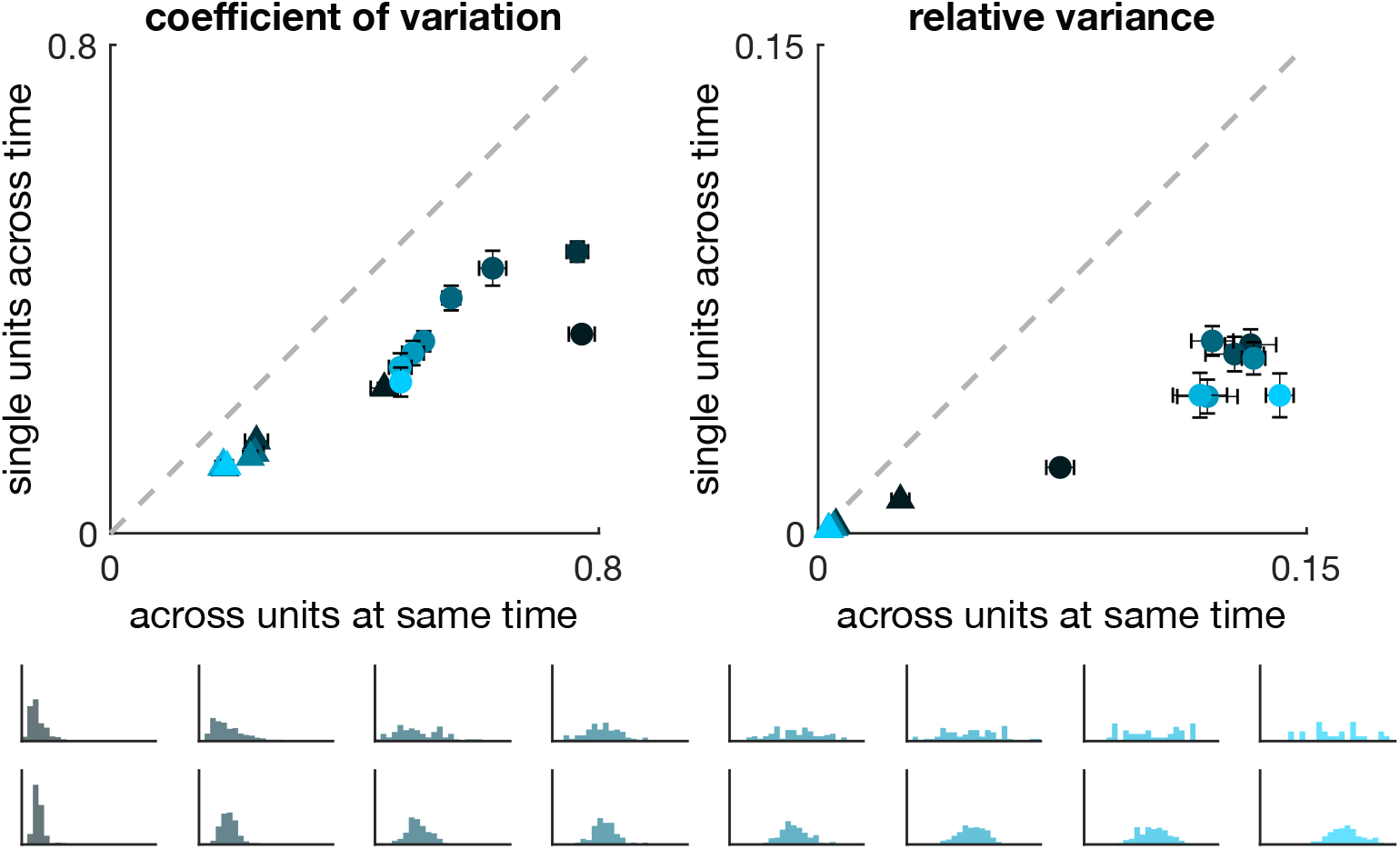
Variability as in Fig. 3 for 4-cell patches.

The sigmoidal intensity correction was used for all experiments. The flattening correction procedure was carried out before experiment start on an area of the plate unoccupied by cells, and was performed for all experiments except the dose response experiments.

### S1.3 Patterning experiments

#### S1.3.1 Global oscillations in score at experiment start

The apparent global oscillations in score before contrast emerges in the *K* = 0.1 case (Fig. 5a) may be a consequence of the fact that cell responses are delayed relative to the frequency with which administered inputs are changed. Intuitively, since all patches begin with similar scores, then during the next imaging period the inputs to all patches will also be roughly the same magnitude. Assuming the scores are initially low, then the lateral inhibition relationship ensures that inputs during the next imaging period will all be similarly high. Cell scores will continue to increase for about two periods following a period of high input, such that an intervening single period of lower input will not drastically affect cell score until about two periods later, when the input has decreased even further due to the continued rise in score from the first high input. A similar effect causes a steep drop in cell scores, restarting the cycle. Imperfect initial conditions and stochasticity ensure that the oscillations eventually decay, allowing for convergence to the contrasting steady state.

#### S1.3.2 Permutation tests to verify contrast

Stochasticity introduces some difference between the mean scores of any two sets of patches regardless of whether they alternate. We conducted statistical tests to verify that the difference in mean between sets of alternating patches exceeded what would be expected for any other arbitrary grouping of patches into two equal-sized sets, i.e., that the contrast arose as a consequence of lateral inhibition separating the patches into two distinct populations of low- and high-scoring cells, rather than due to chance variation in mean scores alone for cells belonging to the same population. In particular, for each experiment we conducted permutation tests against an empirical distribution of *N* = 5000 relabelings, with the null hypothesis that scores for alternating patches were drawn from the same distribution (i.e., the partition into sets of alternating patches is arbitrary). Tests were conducted three times per experiment with results reported in Table S1.4. The contrast in the *K* = 0.1 case was statistically significant in all experiments, while the apparent contrast in the *K* = 1 case was statistically indistinguishable from contrast between random sets of patches. For each of *K* = 0.2 and *K* = 0.3, the majority (2 out of 3) of replicates yielded results consistent with theoretical notions of mono- and bistability at a p-value of 0.05.

#### S1.3.3 Temporal variability in score

Although variability in administered intensity decreased from the first to the last experimental hour in 16-patch experiments, temporal variability in individual patch scores remained nearly the same throughout the experiment (Fig. S1.5a). The result is not surprising if cells were comparably variable regardless of input intensity, in which case convergence of the mean score need not imply reduction in temporal variability. The administered intensity may also appear less variable since it was calculated based on an average over the scores of 4 neighbors, which at a given time step would have a variance 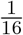 that of an individual patch.

**Figure S1.4:**
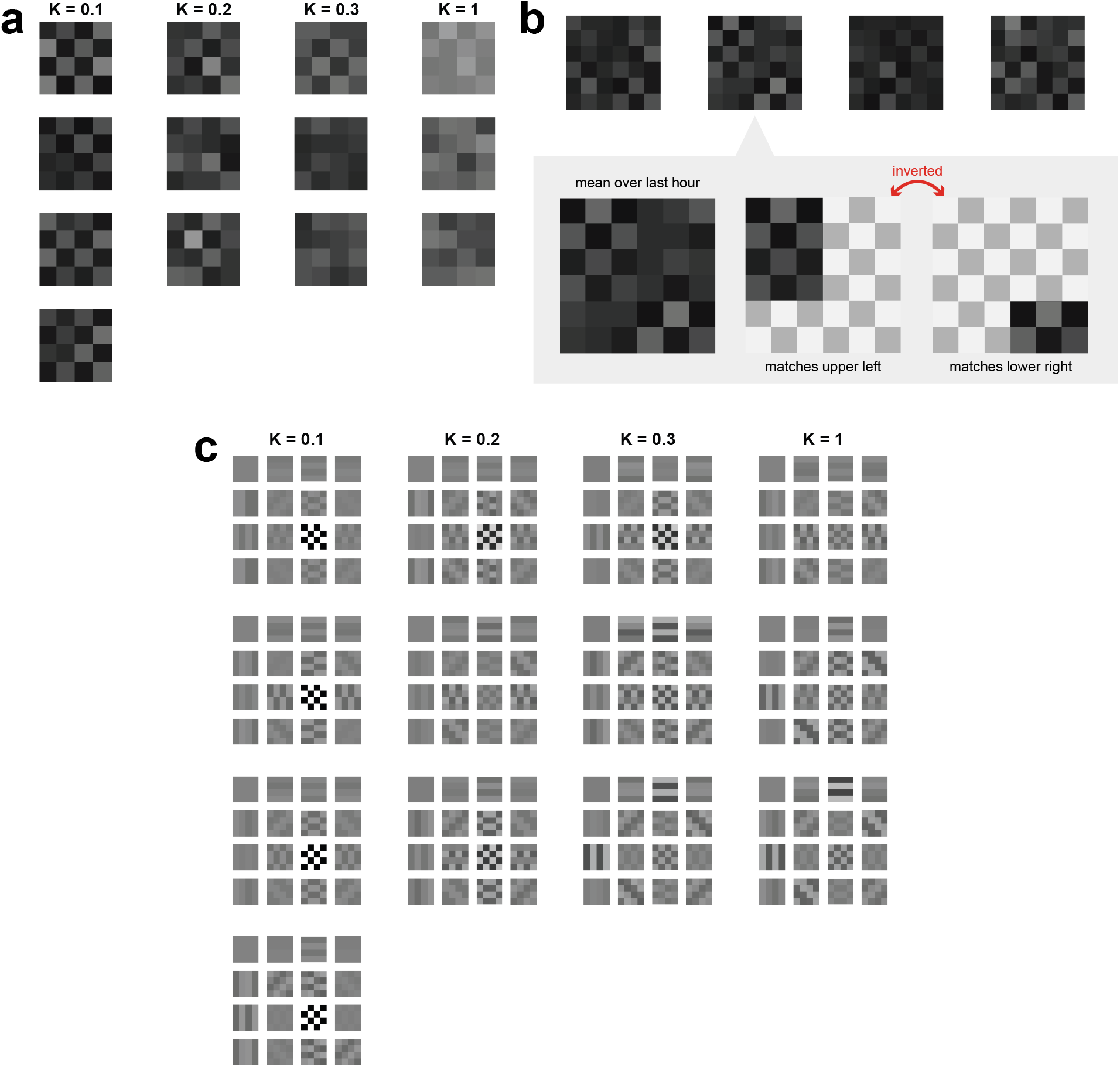
Checkerboards from scores averaged over the last hour for (a) 16-patch and (b) 36-patch (*K* = 0.1) patterning experiments. Although the 36-patch experiments did not achieve global checkerboard patterning, one experiment showed persistent local patterning in which two small checkerboards appeared in opposing corners. The local patterns were inverted relative to each other and did not resolve before the end of the experiment. (c) 2D Fourier transforms conducted on the checkerboard averages from (1) reveal that the greatest weight is given to the highest-contrast spatial mode in all 4 experiments with *K* = 0.1, 2 out of 3 experiments with *K* = 0.2, 1 out of 3 experiments with *K* = 0.3, and none of the 3 experiments with *K* = 1. Pictured are the spectral components (horizontal frequency increasing top to bottom, vertical frequency increasing left to right) that contribute to last-hour checkerboards for each experiment. The intensity of each component indicates its weighting relative to other components in the same experiment. Note also the relatively higher weighting for lower-frequency spatial modes in the *K* = 0.3 and *K* = 1 cases relative to the *K* = 0.1 and *K* = 0.2 cases.

**Table S1.4:** Two-tailed permutation tests for difference in mean score averaged across the last hour between sets of alternating patches, conducted against empirical distributions constructed from N = 5000 relabelings for each experiment individually. The empirical distributions of difference in means between relabeled sets of patches were randomly drawn three times per experiment and the resulting p-value calculated for the alternating patch allocation. Tests were performed against the null hypothesis that scores were drawn from the same distribution for all patches. For *K* = 0.1, the null hypothesis was rejected at a p-value of 0.0002 in all four experiments. For *K* = 1, the null hypothesis failed to be rejected at any reasonable p-value. Results were less clear for values close to the critical point, for which one experiment with *K* = 0.2 (theoretically bistable) failed to reject at any reasonable p-value, and one experiment with *K* = 0.3 (theoretically monostable) rejected at a p-value of 0.02.

| K | trial1 | trial2 | trial3 |
| --- | --- | --- | --- |
| 0.1 | 0 | 0.0002 | 0 |
| 0.1 | 0 | 0 | 0 |
| 0.1 | 0 | 0 | 0 |
| 0.1 | 0 | 0 | 0 |
| 0.2 | 0.0006 | 0.0008 | 0.0008 |
| 0.2 | 0.4422 | 0.4348 | 0.4356 |
| 0.2 | 0.0234 | 0.0226 | 0.017 |
| 0.3 | 0.0158 | 0.0156 | 0.0164 |
| 0.3 | 0.1262 | 0.1282 | 0.1322 |
| 0.3 | 0.146 | 0.1348 | 0.1388 |
| 1 | 0.6516 | 0.6444 | 0.645 |
| 1 | 0.4108 | 0.4116 | 0.4018 |
| 1 | 0.1966 | 0.1942 | 0.2144 |

**Figure S1.5:**
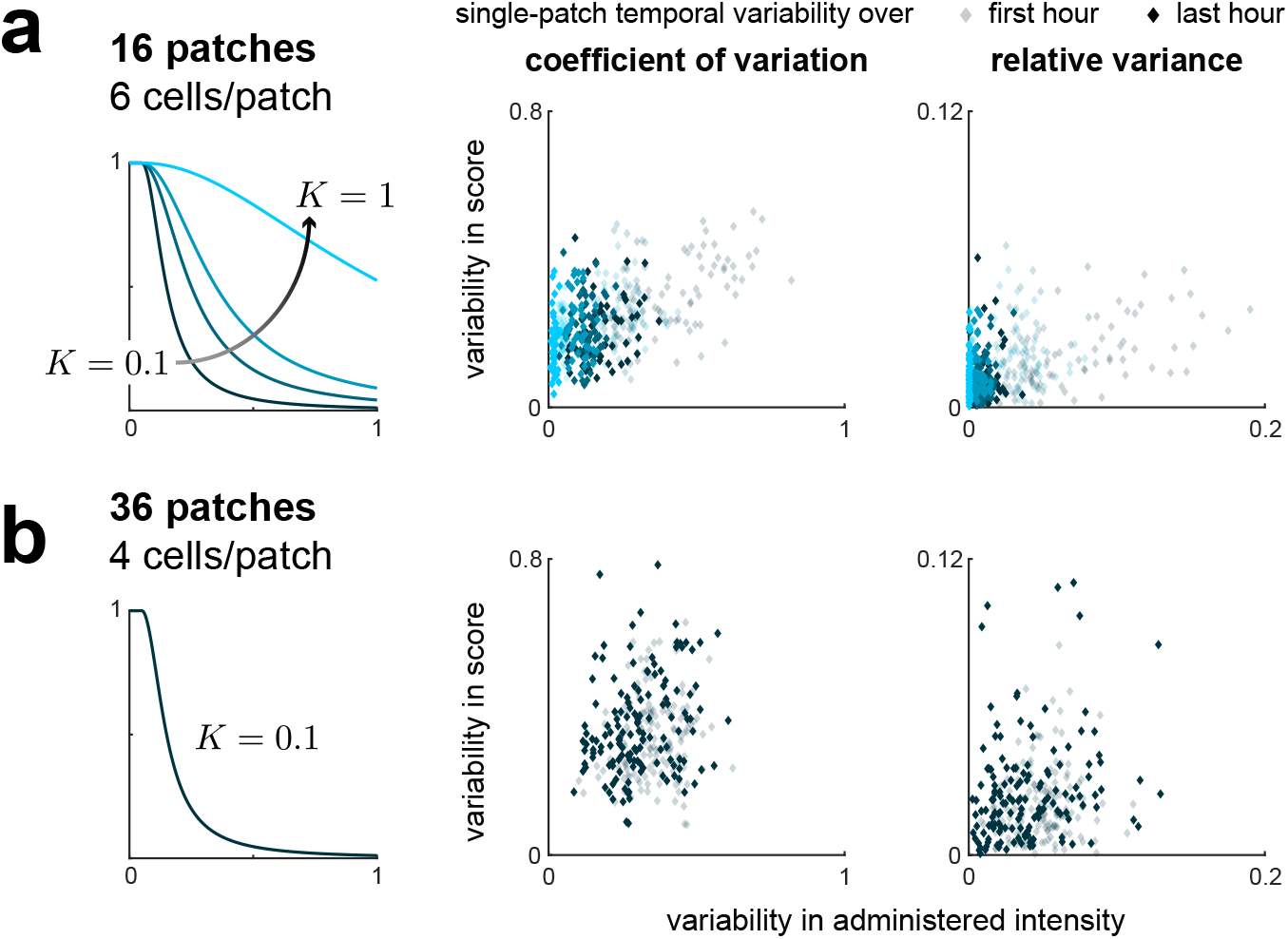
(a) Temporal variability in administered intensity to individual patches decreased from the first to the last experimental hour, while temporal variability in score remained relatively constant. (b) In four experiments with 36 patches and 4 cells per patch, temporal variability in administered intensity and score for individual patches remained equally high throughout the experiment duration (3 hr), suggesting steady state was not reached in that time.

**Figure S1.6:**
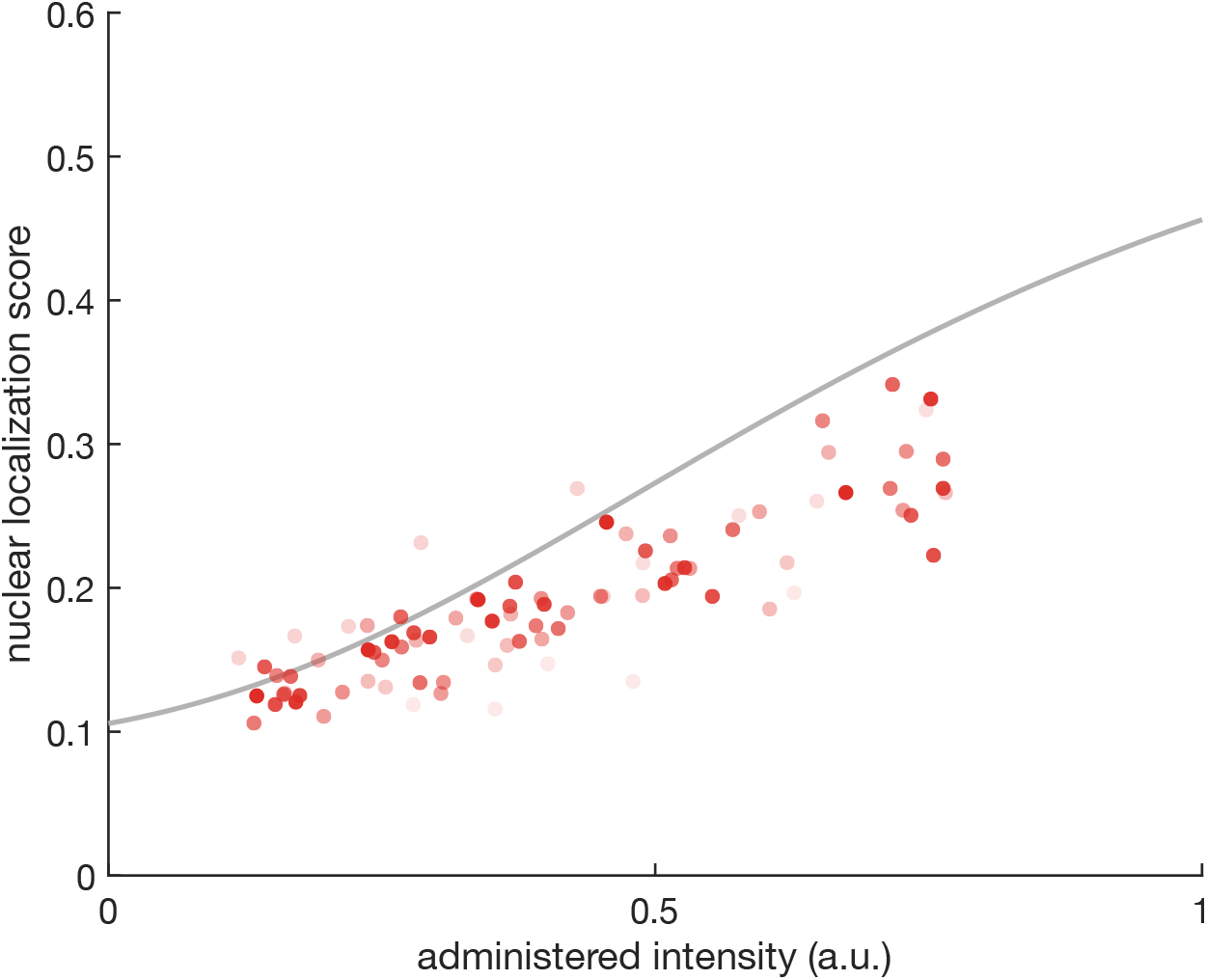
Administered input vs. measured patch score over time in the 36-patch experiments does not show convergence to the steady-state dose response curve in 3 hr, unlike in the 16-patch case (Fig. 5). Points are quantiles across *N* = 4 experiments with *K* = 0.1 for single time points, with more opaque (darker) points corresponding to later times.

**Figure S1.7:**
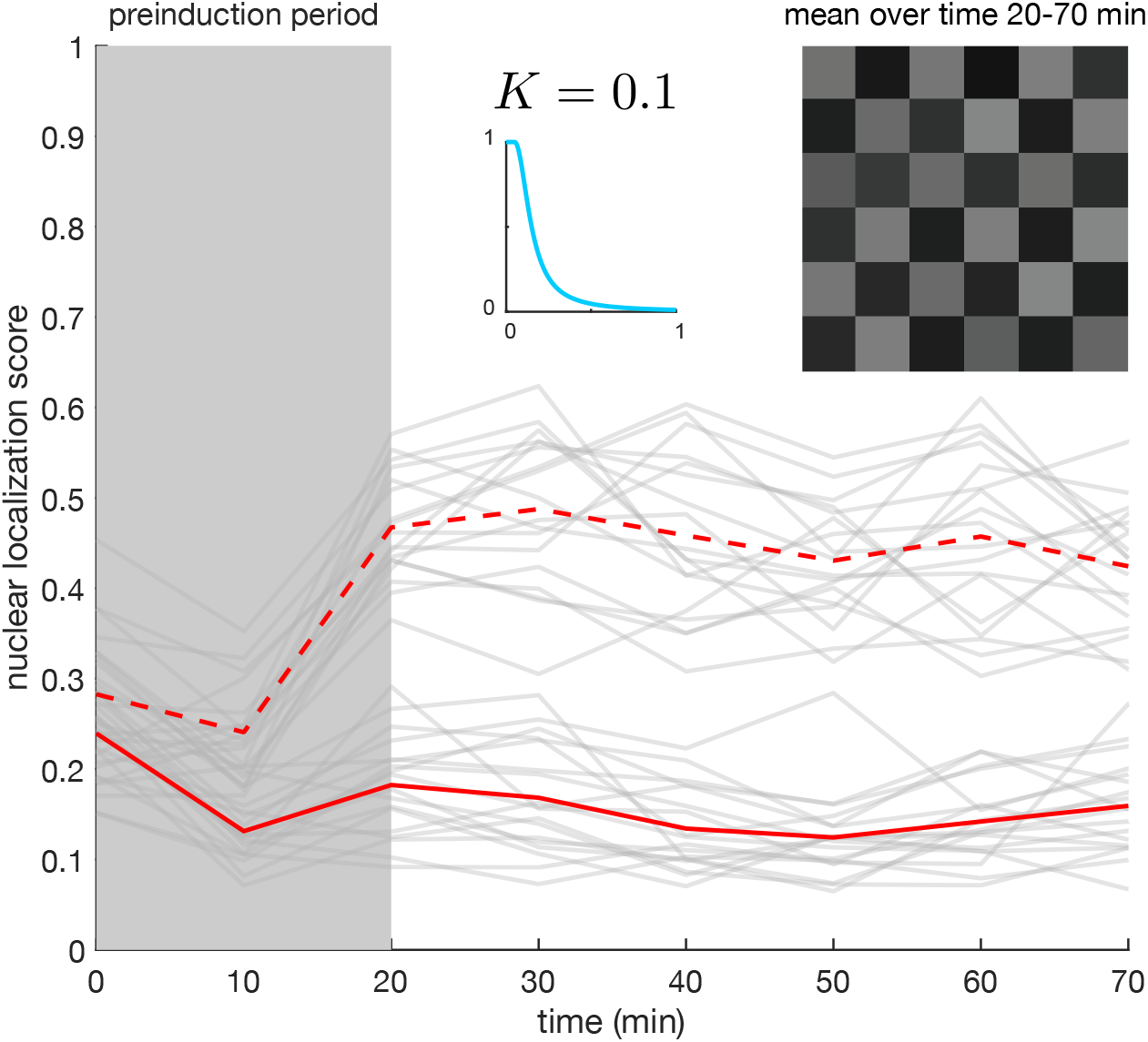
Control experiment verifying persistence of checkerboard pattern in full lateral inhibition system with 36 patchs, 4 cells/patch. Patches were preinduced for 20 min to display a checkerboard pattern. From 20 to 70 min, the system was run in closed-loop with lateral inhibition signaling relation *K* = 0.1. Pictured is the time course for individual patch scores (gray) with averages over sets of alternating patches in red (as in Fig. 4). Board is visualization for individual patch scores averaged over the last 50 min.

